# PIGMENT: A deep learning framework for Porcine Immunohistochemistry seGMENTation

**DOI:** 10.64898/2026.06.18.733245

**Authors:** Pushkar Ambastha, Javid Dadashkarimi, Sai Krishna C. Annavazala, Drew Parker, Ramon Diaz-Arrastia, Hailong Song, Rebecca P Donahue, Douglas H. Smith, Jean-Pierre Dollé, Victoria E. Johnson, John A. Wolf, Ragini Verma

## Abstract

Traumatic brain injury produces widespread axonal damage can be assessed histologically using amyloid precursor protein (APP) immunohistochemistry, which labels injured axonal profiles at cellular resolution [1, 2]. However, quantification of APP pathology remains a major bottleneck: annotation is manual, time-consuming, spatially localized, and variable across raters, limiting scalability and reproducibility. This limitation is particularly important in studies that use histology as a reference for neuroimaging or other tissue-level measurements, where cellular APP pathology must be quantified in a spatial form that can be aligned with imaging abnormalities.

Here, we introduce PIGMENT, an annotation-efficient deep-learning framework for automated segmentation and quantification of APP-positive pathology in porcine white matter histology. PIGMENT uses a compact SegFormer-B0 architecture trained on 525 expert-annotated 512 × 512-pixel tiles from four APP-stained sections across three pigs. Because APP-positive profiles are sparse, fragmented, stain-variable, and morphologically diverse, PIGMENT combines limited expert labels with APP-specific augmentation designed to model variation in APP-positive intensity, size, continuity, fragmentation, and local tissue context.

We evaluated PIGMENT using an instance-level detection rate that measures whether discrete APP-positive components are localized. Across held-out APP-stained data, PIGMENT achieved a mean instance-level detection rate of 0.86. Across the configurations tested, the highest mean detection rate was achieved by a training set that included sections from different animals, suggesting that annotation diversity may be an important factor under limited-label conditions.

By extending limited high-confidence expert annotations into whole-section APP burden maps, PIGMENT provides a scalable framework for characterizing the extent and spatial distribution of traumatic axonal injury. These maps may support future studies that align histological injury burden with imaging-derived measures.

## Introduction

Traumatic brain injury (TBI) results in widespread axonal damage that underlies long-term neurological disability. At the cellular level, traumatic axonal injury can be identified using amyloid precursor protein (APP) immunohistochemistry, a widely used marker of injured axonal profiles with high spatial resolution [3–7]. APP staining provides a detailed map of axonal injury that cannot be obtained through neuroimaging, making it a critical tool for characterizing the extent and spatial distribution of tissue damage.

APP pathology is difficult to quantify because annotation is typically performed manually, requiring expert delineation of APP-positive structures across histological sections [8]. This manual workflow is time-consuming and labor-intensive, and it requires contextual judgment because APP-positive staining must be distinguished from blood vessels, tissue artifacts, and nonspecific background staining that do not represent injured axons. Additional variability can arise because three-dimensional axonal profiles are interpreted from two-dimensional histological sections [9, 10]. These constraints limit the scalability and reproducibility of APP quantification and make it difficult to generate spatially resolved maps of axonal injury burden across large tissue areas.

This limitation becomes a major barrier in studies that aim to use histology as a reference standard for validating neuroimaging. Imaging techniques can detect changes in tissue microstructure, but their interpretation in TBI depends on linking these signals to underlying cellular injury. Establishing this relationship requires quantitative maps of axonal damage that preserve anatomical location and can be aligned with imaging data. Without scalable methods to generate such maps, histology remains limited as a reference for imaging validation.

In addition to the scaling, porcine APP histology presents a difficult image-analysis problem. Staining intensity can vary across batches, scanner settings can affect image appearance, and tissue preparation can introduce artifacts such as folds, tearing, uneven background staining, and section-edge effects [11, 12]. APP-positive structures are multifocal and morphologically diverse, appearing as stereotypical axonal swellings, terminal bulbs, and fusiform profiles [13]. Beyond shape diversity, APP annotation also requires anatomical context, because brown APP-positive immunostaining is interpreted as pathology only when it is consistent with an injured axonal profile, whereas similar brown signal around blood vessels, tissue folds, or nonspecific background staining may introduce false-positive interpretation. Together, these factors make APP segmentation difficult because the model must distinguish true injured axonal profiles from staining variation, tissue artifacts, and APP-like false-positive structures.

These annotation and image-analysis challenges motivate a task-specific automatic segmentation approach. Supervised segmentation models generally benefit from large and diverse annotated datasets, while expert APP annotations are limited and drawn from selected tissue regions [14, 15]. The model must therefore learn from a small set of high-confidence examples while generalizing to new sections, animals, staining conditions, and injury models. Generic augmentation can improve robustness to image-appearance changes such as brightness, contrast, blur, and color variation [16], but it does not necessarily model the APP-specific variation that matters for this task, including fragmented profiles, faint deposits, elongated axonal structures, and APP-like false-positive staining. Similarly, standard segmentation metrics such as Dice coefficient and intersection-over-union are useful for measuring pixel-level agreement, but they can be dominated by larger or geometrically complex APP-positive structures. For burden quantification, the relevant question is also whether individual APP-positive profiles are present and localized, not only how many pixels overlap [17].

Here, we introduce PIGMENT, an annotation-efficient deep-learning framework for automated segmentation and quantification of APP-positive pathology in porcine white-matter histology. PIGMENT uses a compact SegFormer-B0 segmentation model trained from limited expert annotations (Figure 1). Rather than attempting to replace expert judgment, PIGMENT uses high-confidence expert-labeled image regions (tiles) as the foundation for a scalable workflow that extends APP quantification over larger tissue areas. The goal is to use expert annotations as training data to generate reproducible APP burden maps that support downstream anatomical and imaging analyses.

**Figure 1.**
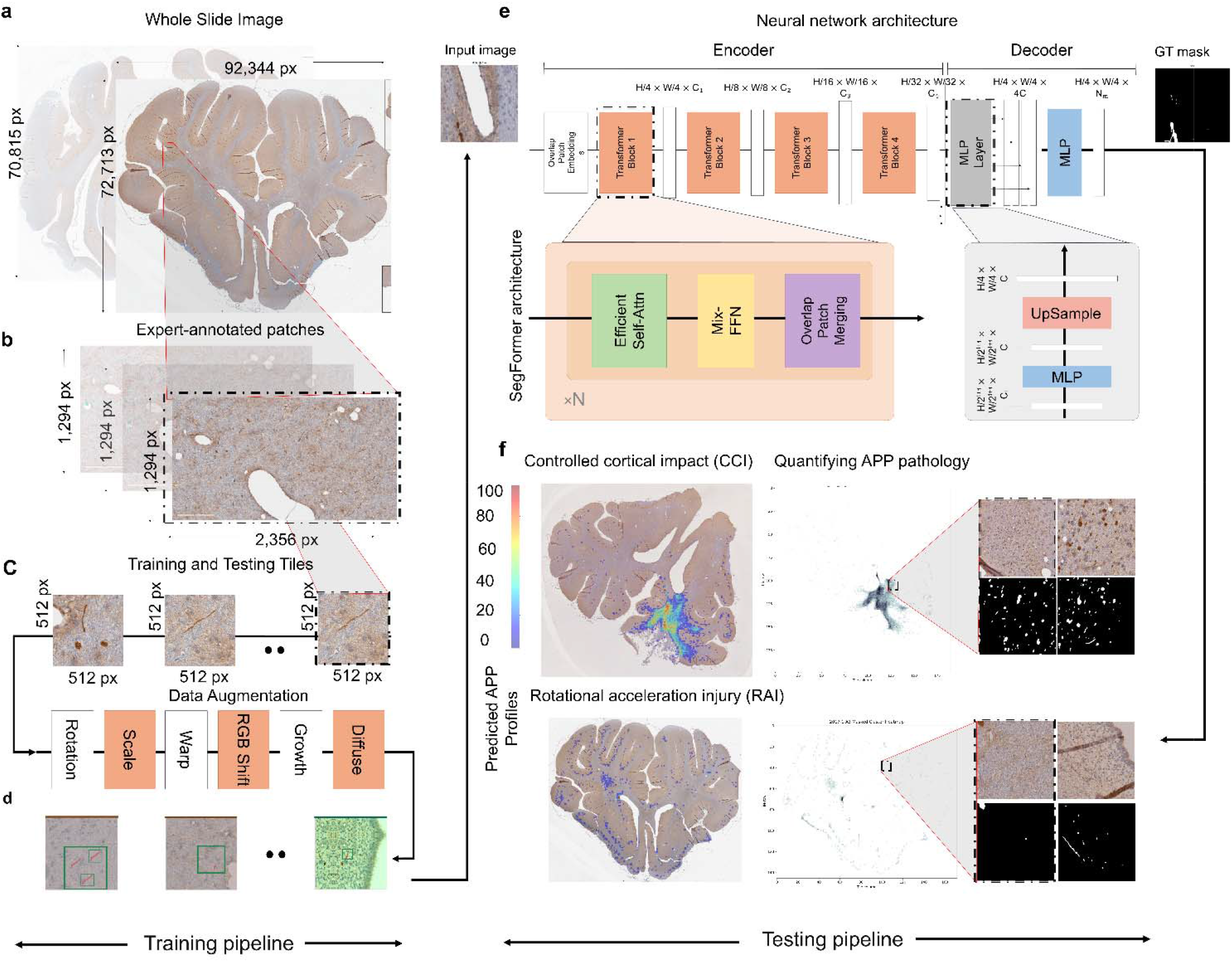
Overview of the PIGMENT workflow. (a) A whole-slide image (WSI) of an APP-stained porcine white-matter section. Each section was first imaged as a full WSI. (b) A trained anatomist manually selected and annotated rectangular patches within the WSI. Patch size varied across annotations and was much smaller than the full section. (c,d) Fixed 512 × 512-pixel tiles were then extracted from within the annotated patches and used as the unit for model training. Example tiles are shown together with representative augmentation operations applied during training. (e) SegFormer-B0 architecture used for APP-positive segmentation. The model takes a 512 × 512 input tile and predicts a corresponding binary APP mask. (f) Testing and deployment pipeline. Model predictions from individual tiles are aggregated to generate section-level spatial maps and heatmap summaries, including examples from rotational acceleration injury and controlled cortical impact-injury tissue.

We designed the workflow around the main requirements of APP histology segmentation. First, because APP-positive profiles vary in appearance and local tissue context, our proposed augmentation strategy varies both image appearance and APP-positive structures while preserving transformed image–mask correspondence. Second, because expert annotations are limited, the sampling strategy tests whether training examples sampled across animals and sections improve held-out performance compared with adding samples from related tissue contexts. Third, because APP burden mapping depends on recovery of individual pathological profiles, the evaluation strategy includes both pixel-overlap metrics and component-level detection [18, 19].

Using 525 expert-annotated 512 × 512-pixel tiles from four APP-stained sections across three pigs, we evaluated whether PIGMENT could generalize across held-out test tiles from each section. We tested whether APP-specific augmentation improves robustness, whether cross-animal and cross-section annotation diversity affects performance, and whether component-level detection provides a biologically meaningful summary at the end.

PIGMENT addresses a key histological bottleneck in large-animal TBI research: the lack of scalable methods for quantifying APP pathology across large tissue areas. By learning from limited expert annotations, PIGMENT generates tissue-level APP maps that retain anatomical location and quantify the distribution of axonal damage. These maps may support future studies that align APP pathology with neuroimaging-derived measures especially in white-matter injury. The present study does not claim to validate neuroimaging biomarkers directly; rather, it establishes the automated histology quantification needed to make such validation studies feasible at larger scale.

## Materials and Methods

### Animals, sections, and annotation

The dataset consisted of four APP-immunostained white-matter sections (D1-D4) from three pigs (P1-P3) in an established rotational acceleration injury (RAI) cohort [20]. Each stained section was scanned at 200× magnification as a whole-slide image. The dataset contained both within- and across-animal variation, where pig P1 contributed section D1, pig P2 contributed sections D2 and D3, and pig P3 contributed section D4.

Three trained investigators manually selected rectangular white-matter regions (patches) within each whole-slide image and annotated APP-positive axonal profiles inside those regions (see Figure 2) [21]. Whole-slide image dimensions ranged from 88,536 to 100,912 pixels in width and from 61,565 to 82,386 pixels in height. Patch dimensions varied across annotations and were smaller than the full whole-slide image; for example, the annotated patch shown in Figure 2 measured approximately 2,356 × 1,294 pixels. Fixed 512 × 512-pixel tiles were then extracted from the annotated patches using a sliding-window procedure and served as the unit for model training and evaluation. This tile size was chosen to provide sufficient local tissue context around APP-positive profiles while remaining compatible with batch-level training on a single GPU. Each tile was paired with a binary mask labeling APP-positive tissue versus APP-negative tissue and background.

**Figure 2.**
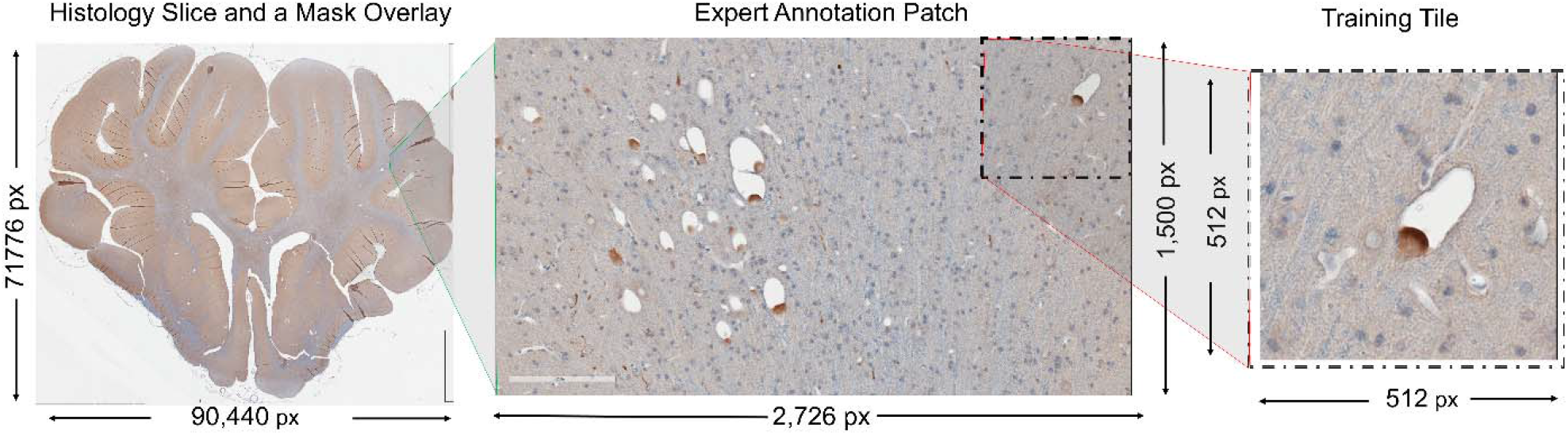
Patch-based expert annotation and tile extraction define the PIGMENT training data. Representative APP-immunostained porcine histology section from dataset D2, illustrating the relationship between the whole-slide image, expert-annotated patch, and fixed-size training tile. The left panel shows the full histology section with the annotated regions. The middle panel shows the corresponding high-resolution annotation patch, where APP-positive axonal profiles were manually labeled by expert annotators. Patch size was constrained by the practical field-of-view and screen-size limitations of the annotation workflow, including Aperio ImageScope and Photoshop. The right panel shows a 512 × 512-pixel tile extracted from the annotated patch using sliding-window sampling. These standardized tiles allow the paired histology image and binary APP mask to fit into GPU memory during training while preserving local cellular-resolution annotation.

**Figure 3.**
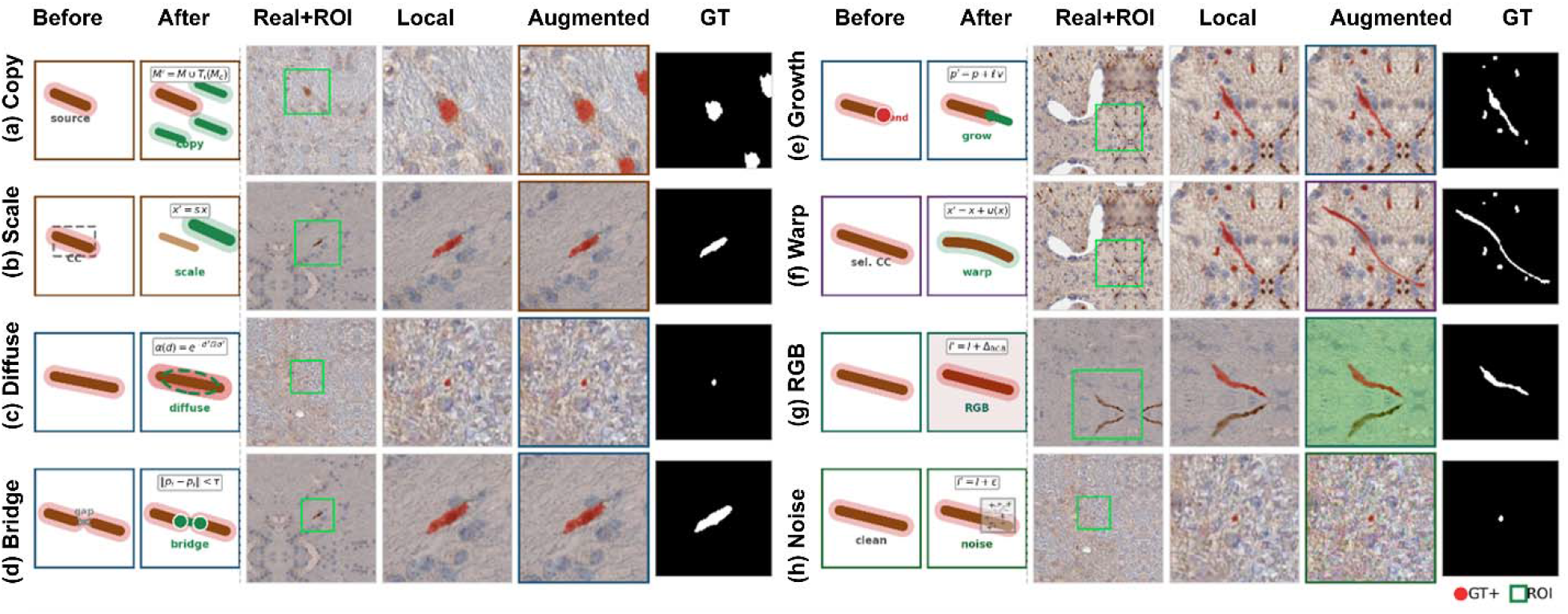
Representative target-aware augmentations applied locally to APP-positive histology regions. Each row shows one transform from the PIGMENT augmentation pipeline: (a) instance copy-paste, (b) component-scale paste, (c) ground-truth neighborhood diffusion, (d) bridge nearby endpoints, (e) growth along the local skeleton, (f) per-component elastic warp, (g) RGB shift, and (h) patch Gaussian noise. Columns show the schematic operation before and after augmentation, the original histology tile with the selected region of interest boxed, the local real crop before augmentation, the locally augmented crop, and the corresponding augmented binary ground-truth mask. The pipeline models both APP-positive morphology and staining variability while preserving image–mask correspondence.

Pixels near patch and tile boundaries were handled conservatively for different reasons. A patch boundary reflects the edge of the anatomist-selected annotation region; APP-positive structures crossing this boundary may be only partially annotated. A tile boundary reflects the edge of the 512 × 512-pixel computational crop used for model training and evaluation; APP-positive structures crossing this boundary may be only partially visible to the model. Therefore boundary-truncated structures below the minimum-size threshold were excluded from both training loss and evaluation, preventing uncertain edge pixels from contributing to model optimization.

The dataset of annotated tiles was split into training (80%) and testing (20%) samples, which are reported in Table 1 for each dataset. In every training configuration, performance was evaluated on held-out tiles from all four sections independently.

**Table 1.** Per-section tile counts in the porcine APP histology dataset, by source pig and train/test split. Three pigs contributed four sections. Pig P2 contributed two sections, whereas P1 and P3 each contributed one section. Each section was split approximately 80% training and 20% testing.

| Dataset | Source pig | Total tiles | Train | Test |
| --- | --- | --- | --- | --- |
| D1 | P1 | 225 | 180 | 45 |
| D2 | P2 | 75 | 60 | 15 |
| D3 | P2 | 90 | 72 | 18 |
| D4 | P3 | 135 | 108 | 27 |
| Total | , | 525 | 420 | 105 |

### Immunohistochemistry

Standard staining protocol was followed, as previously described [20]. Briefly, 8 μm sections were prepared from formalin-fixed paraffin-embedding tissue blocks and mounted on Superfrost microscope slides. Sections were then deparaffinized, followed by rehydration using a series of graded ethanol and water. Next, sections were incubated with 3% hydrogen peroxide to quench endogenous peroxidase activity, then processed for heat-induced antigen retrieval in Tris-EDTA buffer (pH 8.0). Thereafter, sections were blocked with normal serum and incubated with amyloid precursor protein (APP) (Millipore, MAB348, 1:80,000) overnight, followed by incubation with biotinylated secondary antibody and visualization using an avidin-biotin complex and DAB peroxidase substrate kit. Sections were counterstained with hematoxylin for cell nucleus and digitally scanned with a 20x objective on an Aperio AT2 slide scanner.

### Architecture

The proposed PIGMENT, technically, maps each tile to a pixel-wise probability map of APP positivity. For fair comparisons, we fixed our architecture design to SegFormer-B0, a compact hierarchical transformer segmentation model with a Mix Transformer encoder and lightweight decoder [22]. As such, the encoder produces multiscale feature maps that are fused by the decoder to generate a full-resolution output mask. Another reason to choose SegFormer-B0 was because it has relatively few parameters compared with larger segmentation backbones, making it suitable for a limited annotated dataset, while still combining fine and coarse image features during prediction. The utilized model contains approximately 3.7 million parameters and was initialized with publicly available ImageNet-1k pretrained weights before fine-tuning on APP tiles.

The segmentation head produced a single-channel probability map with the same spatial dimensions as the input tile [16]. During inference, the probability map was thresholded at 0.5 to obtain a binary APP-positive prediction. Full architecture details are provided in the Supplementary Information.

### Training objective and optimization

Models were trained using a pixel-wise Dice loss computed only over annotated, valid pixels. To reduce edge effect, boundary-truncated regions were excluded from the loss so that partially annotated APP-positive structures did not influence optimization. To further reduce the effect of uncertain labels, we used an exponential moving average teacher model during training [23]. At each iteration, regions with strong disagreement between the teacher prediction and the manual annotation were excluded from the loss for that iteration only. This label-noise filtering step was applied only during training and did not change the expert annotations used for evaluation. Models were optimized using AdamW [18] with learning rate of 1 × 10^−^□, β_1_ = 0.9, β_2_ = 0.999, and weight decay 0.01. Training used a batch size 8. Dropout was applied to encoder hidden states (p = 0.1), attention weights (p = 0.1), and the segmentation head (p = 0.3). Training a single model on a 24-GB GPU required less than two hours on this dataset.

### APP-specific augmentation pipeline

Augmentation was applied on the fly during training using python’s generator function. This allows us to feed our training with as many variations as needed for generalization. Spatial transformations were applied jointly to the image and mask, whereas photometric transformations were applied only to the image. Basically, the proposed pipeline was designed to expose the model to variation in both tissue appearance and APP-positive profile morphology. We grouped these transformation functions into five classes: spatial, photometric, copy-paste, morphological connectivity, and tissue-level transforms.

Spatial transforms included horizontal and vertical flips, rotations, affine perturbations, and elastic deformation of elongated APP-positive components. Photometric transforms included brightness and contrast shifts, color jitter, Gaussian blur, motion blur, sensor noise, compression artifacts, global RGB shifts, local RGB shifts, and patch-level noise. Copy-paste transforms randomly inserted annotated APP-positive connected components into nearby training tiles while preserving the component mask and their local appearance. Morphological connectivity transforms bridged nearby fragmented components and extended elongated APP-positive profiles along their local orientation. Tissue-level transforms modified background and weak APP-like staining patterns, including white-tissue dropout, Gaussian-mixture-model-based perturbation of pixels with color values similar to annotated APP-positive regions, and local smoothing around annotated APP profiles. Full parameter ranges and implementation details for each transform are provided in the Supplementary Methods.

All transforms preserved image–mask correspondence. The complete list of transforms, probabilities, and parameter ranges is provided in Table 2 and in Supplementary Information.

**Table 2.** Target-aware augmentation pipeline used for PIGMENT training. Transform classes were grouped into geometric, tissue-level, photometric, copy–paste, and mask-guided operations. Spatial transforms were applied jointly to the image and binary APP mask, whereas photometric transforms, marked by †, were applied to the image only or were mask-guided while preserving image–mask correspondence. Here *D* denotes tile width, *M*_*r*_ denotes a sampled regional mask, *b* denotes replacement background intensity, *A*_min_ denotes minimum connected-component area, and *U*(·)denotes a uniform distribution. The dagger (†) marks image-only augmentations that alter staining appearance or local texture while preserving the corresponding binary ground-truth mask. Probabilities denote independent application probabilities within the training augmentation pipeline.

| Transform | Category | Probability | Key parameters / formulation |
| --- | --- | --- | --- |
| Horizontal flip | Geometric | 0.50 | $x' = F_h(x)$ |
| Vertical flip | Geometric | 0.20 | $x' = F_v(x)$ |
| Random 90° rotation | Geometric | 0.30 | $k \sim U[24]; x' = R_{\{90k\}}(x)$ |
| Affine transform | Geometric | 0.40 | $\theta \sim U(-18^\circ, 18^\circ); s \sim U(0.96, 1.06); t \sim U([-0.06D, +0.06D])$ |
| White tissue dropout | Tissue-level | 0.25 | Region-level background replacement:<br>$x' = x \odot (1 - M_r) + b \cdot M_r$ |
| GMM label promotion | Tissue-level | 0.35 | $K = 4; r \leq 4500; n_{region} \leq 3; p_{seed} = 0.80; p_{ignore} = 0.75$ |
| Color jitter block † | Photometric | 0.45 | $x' = T_j(x), T_j \sim OneOf(\tau); \tau$ includes brightness/contrast, ColorJitter, Gaussian blur, motion blur, and ISO noise |
| Global RGB shift † | Photometric | 0.35 | $\Delta R, \Delta B \sim U(-8, 8); \Delta G \sim U(-6, 6)$, applied in the $[0, 255]$ intensity range |
| JPEG compression † | Photometric | 0.20 | $q \sim U(85, 98); x' = \text{JPEG}(x; q)$ |
| Multi-instance copy–paste | Copy–paste | 0.92 | $n \sim U\{5, \dots, 20\}; s_k \sim U(0.90, 1.25); \theta_k \sim U(-8^\circ, 8^\circ); \text{feather} = 2 \text{ px}$ |
| Component-scale paste | Copy–paste | 0.20 | $s \sim U(0.95, 1.90); A_{\min} \geq 140 \text{ px}^2; \text{context radius} = 14 \text{ px}$ |
| GT neighborhood diffusion † | Tissue-level / mask-guided photometric | 0.35 | $r \sim U(4, 10); \alpha \sim U(0.12, 0.28); \tau \sim U(0.75, 0.90)$ |

### Evaluation metrics, and post-processing

PIGMENT generated a probability map for each 512 × 512-pixel tile. Probability maps were thresholded at 0.5 to produce binary APP-positive predictions. All post-processing and metric computation were performed identically for PIGMENT and for the baseline methods.

Before evaluation, we applied artifact and boundary-exclusion rules to exclude potential common artifacts due to tissue preparation and staining, including folding from tissue mounting, tears from sectioning, bubbles from cover slipping, and edge effects. Predicted connected components were excluded if they met both of the following criteria: area ≥ *v* pixels and elongation ratio ≥ *τ*. We empirically validated these parameters to be *v* = 2300*px, τ* = 3.0. Elongation ratio was defined as the ratio of the major-axis length to the minor-axis length of the fitted component ellipse. These thresholds were selected from the training annotations and were used to remove structures that were inconsistent with the annotated APP-positive profiles in this dataset.

The instance-level detection rate (DR) is computed over valid ground-truth components after closing G with a disk of radius r = 2 and dilating P by r′ = 1:

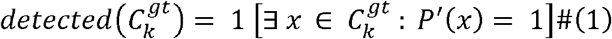

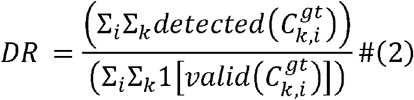

For other two metrics, let G′ and P′ denote the foreground pixel sets of the (masked) ground-truth and predicted binary tile, respectively, and let 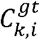 be the set of valid ground-truth connected components after the boundary-aware ignore step. The two pixel-overlap metrics are:

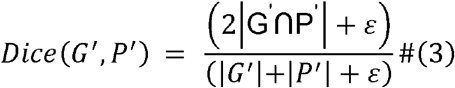

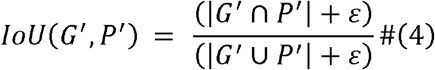

Predicted components smaller than *v*=25 pixels were removed, and ground-truth components smaller than *v*=25 pixels were not included in component-level detection analysis yet everything in ground truth is considered in global detection result. This minimum-size threshold was applied to reduce edge effect [25].

We report instance-level detection rates as the primary metric. To allow minor boundary uncertainty and to fill unexpected holes, the predicted mask was dilated and then eroded by one pixel before overlapping testing. A valid ground-truth component was counted as detected if any pixel of the dilated prediction overlapped that component.

Dice coefficient and intersection-over-union were reported as secondary pixel-overlap metrics. These metrics were computed after applying the same ignore-mask, artifact-removal, and minimum-size rules described above. Full mathematical definitions of detection rate, Dice, IoU, and all post-processing operations are provided in the Supplementary Information.

### Baseline comparisons

We compared PIGMENT with two general-purpose segmentation baselines: UniverSeg, a few-shot segmentation method that uses support image–mask pairs at inference [26], and MicroSAM, a Segment Anything-based microscopy segmentation framework which is inherently designed to segment cell nucli [27]. These baselines were selected to represent two practical alternatives to training a task-specific APP model: support-based segmentation from a small number of examples and zero-shot segmentation using a foundation-model-derived microscopy tool.

UniverSeg was evaluated using the same training-section configurations as PIGMENT. For each configuration, support examples were drawn exclusively from the corresponding training tiles. Training tiles were padded to 576 × 576 pixels, from which 128 × 128 support windows containing the target structure were extracted, matching the input size UniverSeg is designed for. At inference, each held-out 512 × 512 test tile was padded to 576 × 576 and segmented using overlapping 128 × 128 sliding windows with stride 64 to reduce edge effects. Window-level predictions were averaged across overlapping regions, then thresholded to produce a full-tile binary mask.

MicroSAM was evaluated without APP-specific fine-tuning using its automatic mask-generation workflow. Inference was performed independently on each held out 512 × 512 test tile. Candidate masks generated by MicroSAM were combined into a binary prediction map and then evaluated using the same post-processing, ignore-mask handling, and metric pipeline as PIGMENT.

These baselines were included to compare PIGMENT with general-purpose segmentation strategies under the same evaluation framework. They were not intended to exhaustively benchmark all possible prompt-engineered, fine-tuned, or task-specific adaptations of foundation segmentation models.

### Training configuration comparison

To understand what drives generalization under limited annotation, we defined four training configurations that vary the composition of the training set rather than the model or training procedure. The simplest was D1 alone — a single densely annotated section from pig P1, giving 180 training tiles. We used this as our baseline because it represents the most realistic starting point: one animal, one section, the minimum viable annotation effort. From there, we asked whether adding more tiles from a second animal (D1+D2+D3, 312 tiles) or scaling up to everything available (D1+D2+D3+D4, 420 tiles across all three pigs) would meaningfully improve performance. We also tested a fourth configuration (D1+D4) that skips the two additional P2 sections entirely and instead pairs one section from P1 with one from P3, yielding 288 tiles drawn from two different animals. This last configuration was specifically designed to ask whether cross-animal diversity matters more than raw tile count.

To keep the comparison clean, we held everything else constant including architecture, augmentation, training objective, and all hyperparameters, and varied only what went into the training set. Every configuration was then evaluated on held-out test tiles from all four sections independently.

## Code and data availability

### Independent controlled cortical impact sample

In addition to the RAI sections used for model development and quantitative evaluation, we examined one independent controlled cortical impact (CCI) section as a qualitative generalization test. This sample was not included in model training, validation, or hyperparameter selection. Because exhaustive expert annotation was not available for this section, it was not used for formal quantitative external validation. Instead, PIGMENT was applied to the section to assess whether the model could localize APP-positive profiles in visibly affected tissue and generate an APP burden map in an injury setting outside the RAI development cohort.

### Source Code

Source code, the augmentation pipeline, trained model checkpoints along with apptainer container are publicly available at https://github.com/dadashkarimi/pigment.

## Results

### Evaluation across training configurations

Across the four training configurations, weighted mean detection rate ranged from 0.803 to 0.850, with smaller variation in Dice (0.675–0.695) and IoU (0.562–0.578; Table 3.). The baseline configuration (D1) achieved a weighted mean DR of 0.810. Adding within-animal sections from a second pig (D1 + D2 + D3) did not improve overall DR (0.806) despite increasing training volume from 180 to 312 tiles. Incorporating all available sections (D1 + D2 + D3 + D4, 420 tiles) similarly did not improve over D1. The D1 + D4 configuration, which drew one section from each of two different animals, achieved the highest weighted mean DR (0.850), Dice (0.695), and per-section DR across all four held-out sections (0.92, 0.86, 0.75, 0.80).

**Table 3.**
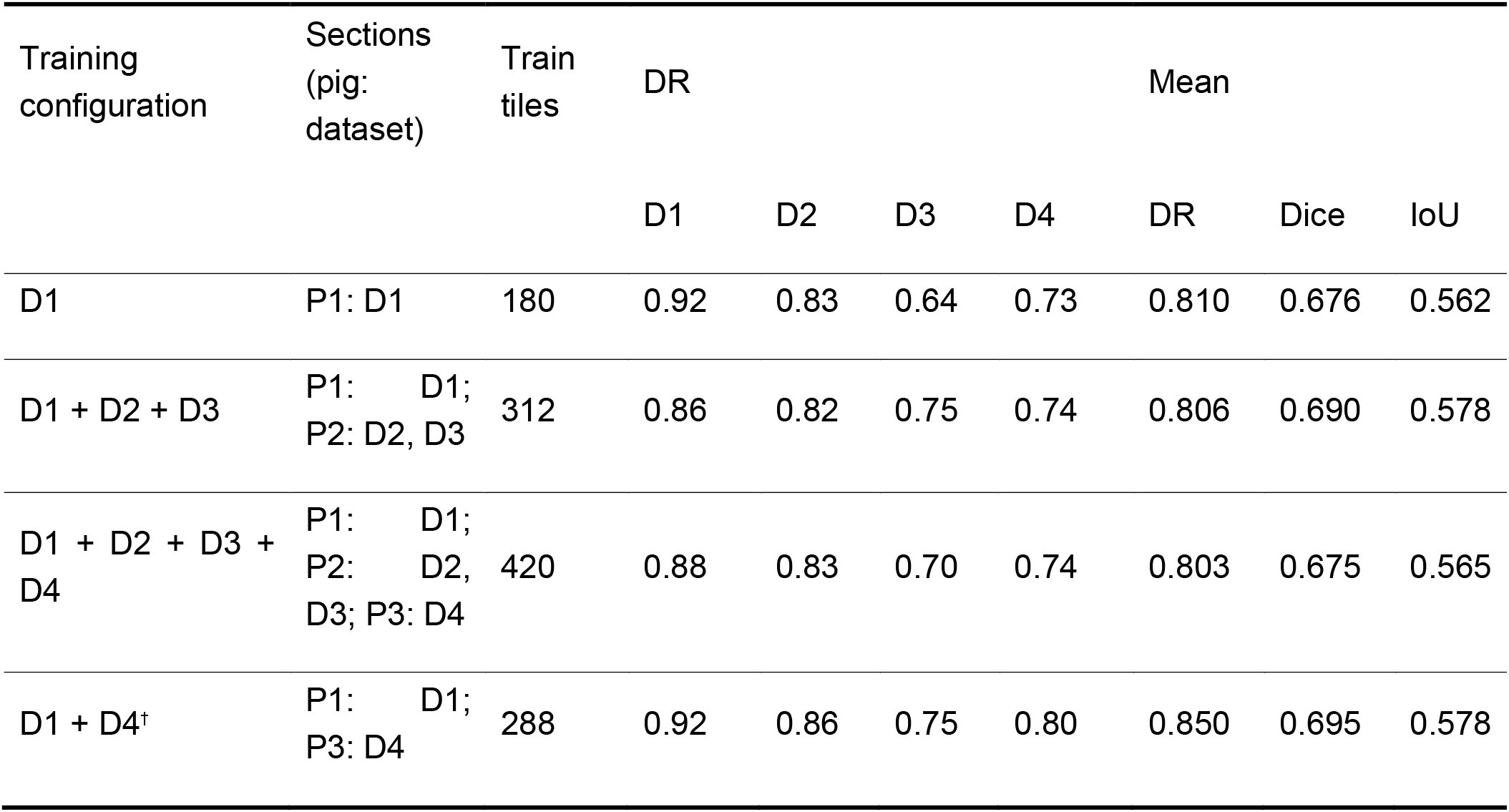
Effect of training-set composition on generalization of PIGMENT. Four training configurations were evaluated: D1 only, D1 + D2 + D3, D1 + D2 + D3 + D4, and D1 + D4. Each model was trained on 80% of the tiles from the selected training sections and evaluated on held-out test tiles from all four sections. Per-section detection rate (DR) is reported for each held-out test section. Mean DR, Dice, and IoU summarize performance across all held-out test tiles (n = 105). DR is an instance-level metric in which a ground-truth APP-positive connected component is counted as detected when any pixel of the dilated prediction overlaps it. Dice and IoU are pixel-overlap metrics. All metrics were computed after artifact-exclusion and boundary-ignore post-processing. Bold values indicate the best mean performance. †Best-performing configuration.

| Training configuration | Sections (pig: dataset) | Train tiles | DR |  |  |  | Mean |  |  |
| --- | --- | --- | --- | --- | --- | --- | --- | --- | --- |
|  |  |  | D1 | D2 | D3 | D4 | DR | Dice | IoU |
| D1 | P1: D1 | 180 | 0.92 | 0.83 | 0.64 | 0.73 | 0.810 | 0.676 | 0.562 |
| D1 + D2 + D3 | P1: D1; P2: D2, D3 | 312 | 0.86 | 0.82 | 0.75 | 0.74 | 0.806 | 0.690 | 0.578 |
| D1 + D2 + D3 + D4 | P1: D1; P2: D2, D3; P3: D4 | 420 | 0.88 | 0.83 | 0.70 | 0.74 | 0.803 | 0.675 | 0.565 |
| D1 + D4† | P1: D1; P3: D4 | 288 | 0.92 | 0.86 | 0.75 | 0.80 | 0.850 | 0.695 | 0.578 |

### Comparison of component-level and pixel-overlap metrics

Representative held-out tiles illustrate the relationship between detection rate and pixel-overlap metrics (Figure 4). Samples 2 and 3 achieved complete component-level detection (DR = 1.00) with moderate Dice values (0.403 and 0.598), indicating that PIGMENT localized the annotated APP-positive profiles despite imprecise boundary agreement. Sample 1 showed partial component recovery (DR = 0.75, Dice = 0.648). Sample 4 demonstrated a similar pattern of partial recovery (DR = 0.67, Dice = 0.479). Sample 5 was a complete failure case (DR = —, Dice = 0.000), where no APP-positive structures were annotated in the ground truth and no predictions were made. These examples confirm that detection rate and Dice capture complementary aspects of performance: boundary disagreement and sparse fragmented profiles lower Dice without necessarily preventing component localization.

**Figure 4.**
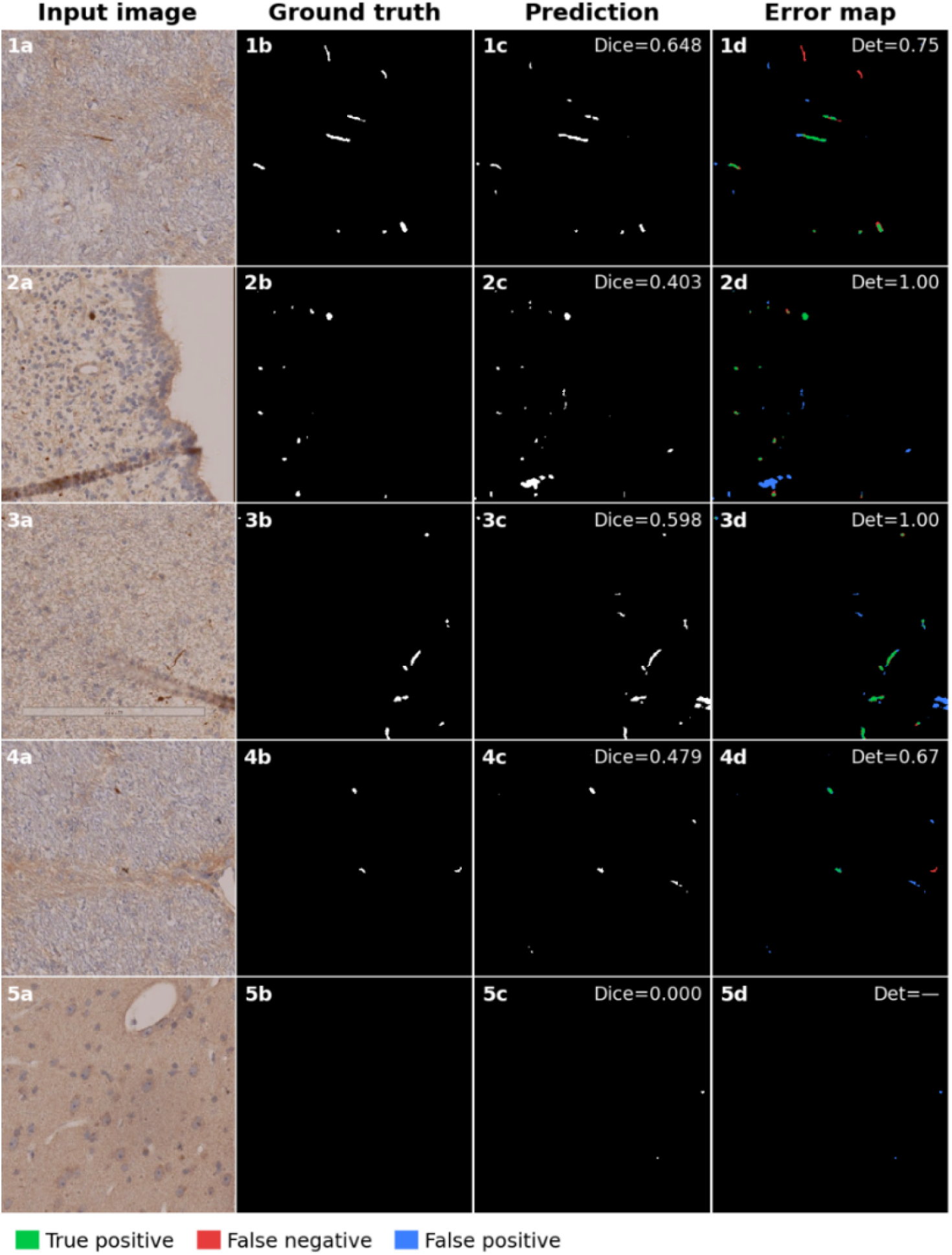
Representative qualitative cases illustrating PIGMENT behavior across held-out APP histology tiles. Each row shows the input histology tile, expert ground-truth mask, PIGMENT prediction, and color-coded error map. Error maps are evaluated at the pixel level and show true positives in green, false negatives in red, and false positives in blue. Dice and component-level detection rate are provided for each example. These cases illustrate why pixel-overlap metrics and detection rate capture different aspects of performance: PIGMENT can localize APP-positive components even when boundary disagreement, sparse fragmented profiles, or low-contrast staining lowers Dice. Samples with detection rate of 1.00 demonstrate successful component localization despite moderate Dice, whereas lower-detection examples highlight missed small or faint APP-positive fragments as a dominant failure mode.

### Comparison with general-purpose segmentation baselines

PIGMENT produced substantially higher Dice values than both baselines across all five representative samples (Figure 5). PIGMENT Dice ranged from 0.59 to 1.00 across samples, compared with 0.00 to 0.34 for UniVerSeg and 0.00 for MicroSAM in all cases. UniVerSeg generated dense false-positive predictions across background tissue, producing near-zero Dice despite visually active output masks. MicroSAM failed to recover APP-positive structures in every sample, consistent with zero or near-zero Dice across the panel. On Sample 4, PIGMENT localized a small APP-positive fragment that both baselines missed entirely. These qualitative patterns are consistent with the quantitative summary: averaged across all evaluated configurations and held-out sections, PIGMENT reached a mean DR of approximately 0.85 and Dice of approximately 0.74, compared with approximately 0.51 DR and 0.13 Dice for UniVerSeg, and near-zero performance for MicroSAM on both metrics.

**Figure 5.**
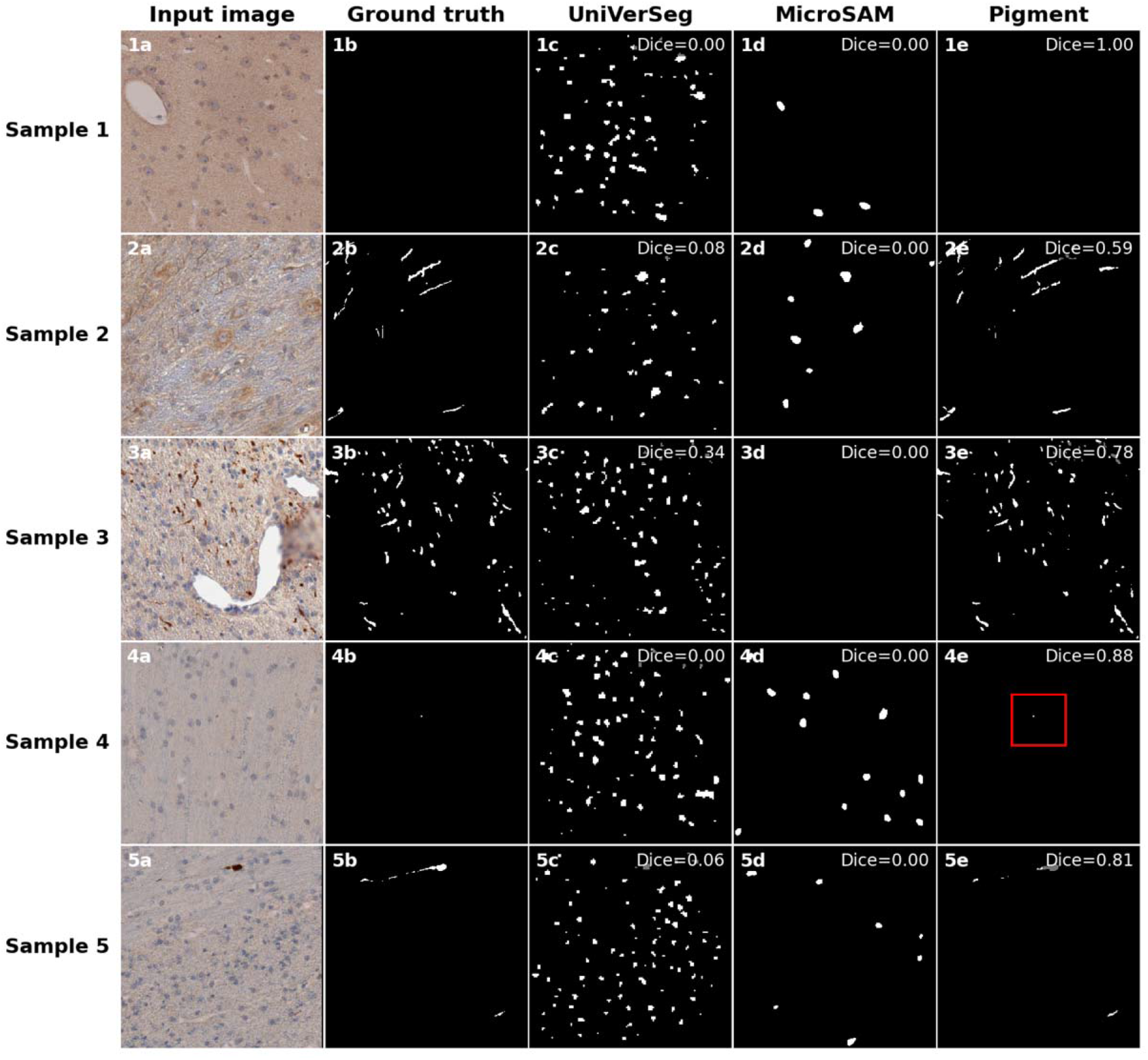
Qualitative comparison of PIGMENT against UniverSeg and MicroSAM on representative held-out APP histology tiles. Columns: input tile, expert ground-truth binary mask, UniverSeg prediction, MicroSAM prediction, PIGMENT prediction. Per-tile Dice values are shown in each panel. Across these cases, PIGMENT recovers the annotated APP-positive components consistently; UniverSeg over-segments background tissue with scattered false positives; MicroSAM under-segments and frequently misses APP-positive structures entirely. The red box on Sample 4 highlights a single small APP fragment that PIGMENT correctly localized while both baselines missed.

### Effect of spatial augmentations

To assess whether spatial augmentation adds value beyond intensity normalization alone, we compared the full-augmentation model against a baseline that used histogram matching but no spatial transformations. The two models performed similarly overall, with full augmentation showing small positive trends on three of the four test sections (Figure 6). The clearest gains appeared on D4 and D3, where full augmentation improved Dice by +0.043 and +0.036, IoU by +0.025 and +0.030, and detection rate by +0.049 and +0.042 respectively. D2 followed the same direction, with improvements of +0.026 Dice, +0.022 IoU, and +0.036 detection rate. D1 was the exception, showing near-zero or marginally negative differences across all three metrics. When pooled across all sections, the overall changes were modest: +0.013 Dice, +0.008 IoU, and +0.022 detection rate.

**Figure 6.**
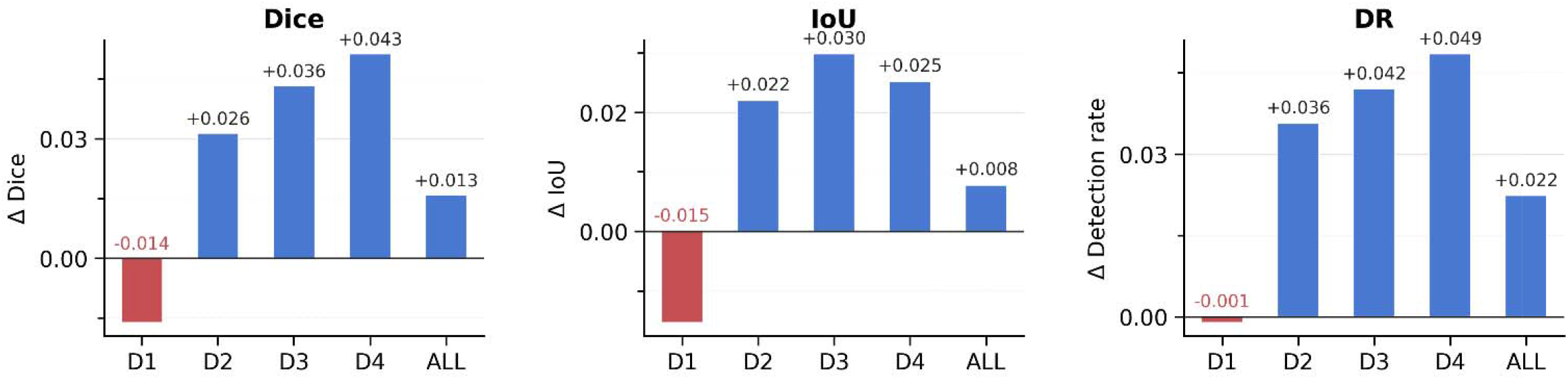
Relative improvement of spatial augmentations compared to histogram normalization baseline, both trained on D1+D4.

### Qualitative evaluation in an independent CCI sample

We applied PIGMENT to an APP-stained section from an independent controlled cortical impact (CCI) injury sample that was not used for model development (Figure 7). Because exhaustive expert annotation was not available, we treated this experiment as a qualitative test rather than a formal external validation benchmark. PIGMENT detected APP-positive profiles in visibly affected tissue and generated a tissue-level map of predicted APP burden. Qualitative review by an experienced neuroanatomist indicated that the predictions captured some APP-positive structures, although certain APP-positive swellings were not detected and positive predictions were also observed in cortical regions where pathology was not expected. Consequently, this experiment provides preliminary evidence that PIGMENT can localize candidate APP-positive regions in an independent CCI sample, but independently annotated external sections or representative test regions will be required for rigorous quantitative assessment of model performance and generalizability.

**Figure 7.**
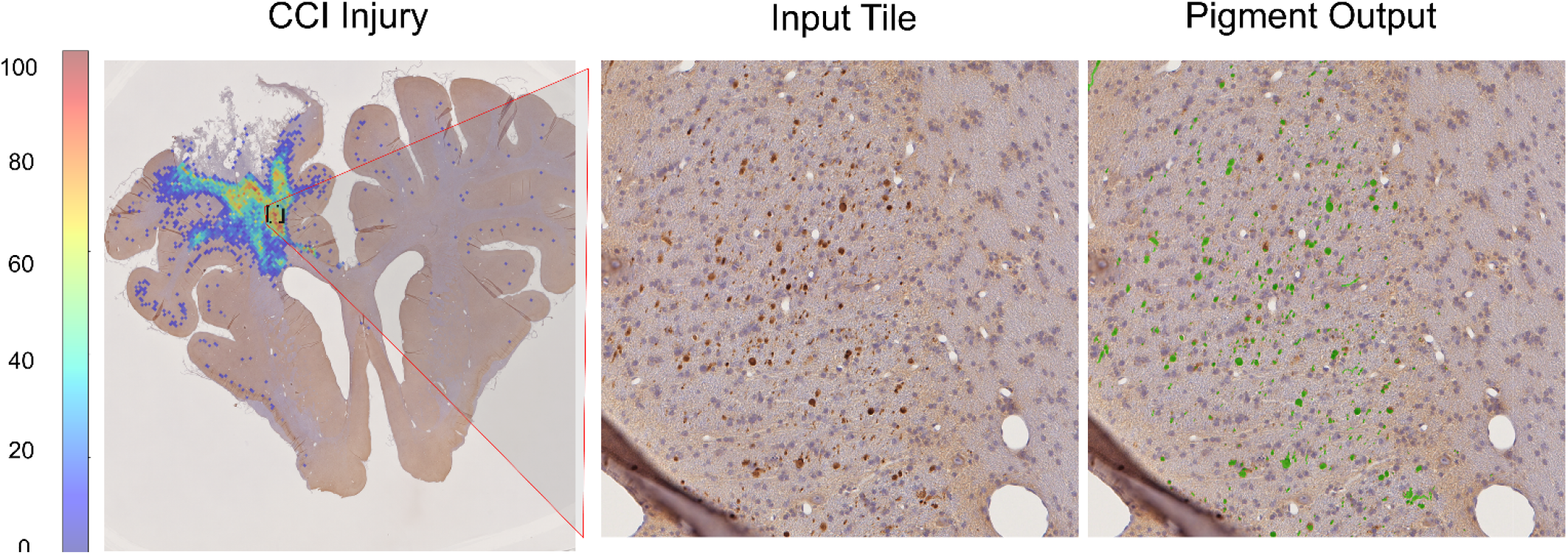
Qualitative evaluation of PIGMENT on an independent controlled cortical impact (CCI) porcine histology sample not included in training. Left: whole-section spatial summary of predicted APP burden, shown as a cluster-intensity map where warmer colors indicate higher local density of predicted APP-positive structures. Middle: representative input tile from the same sample, showing brown APP-positive immunostaining. Right: PIGMENT output overlaid on the input tile, with predicted APP-positive regions shown in green. PIGMENT identifies the focal region of APP-positive pathology as well as scattered deposits in surrounding tissue.

## Discussion and Conclusions

We developed PIGMENT, a deep learning framework for automated segmentation and quantification of APP-positive axonal pathology in porcine brain white matter. Several features of APP immunohistochemistry make this a challenging problem for automation. For example, expert annotations are often limited, labeling is concentrated in selected tissue patches rather than distributed across entire sections, and identifying APP-positive axonal profiles requires contextual anatomical judgment. The target structures are also morphologically diverse, ranging from punctate deposits to fragmented accumulations and elongated curvilinear profiles, with staining intensity that may vary across sections and batches. These features make standard pixel-overlap metrics alone insufficient for evaluating model performance. For quantifying axonal injury burden, recovering individual pathological profiles can be as important as reproducing every positive pixel.

To address these challenges, we developed PIGMENT as a task-specific computational framework rather than relying on segmentation architecture alone. The distinguishing contribution of PIGMENT is the integration of limited high-confidence expert annotations, histogram normalization, APP-specific morphology and appearance augmentation, boundary-aware training, component-level evaluation, and reconstruction of tile-level predictions into spatial tissue maps. This end-to-end design addresses the full workflow from sparse microscopic annotation to scalable tissue-level quantification. Using a compact SegFormer-B0 model trained on 525 expert-annotated tiles from four sections across three pigs, PIGMENT achieved strong component-level detection across held-out sections and generated section-level maps of APP-positive axonal pathology.

We report instance-level detection rate as the primary metric because it reflects how PIGMENT is intended to be used downstream. The goal is to generate tissue-level measurements of axonal injury that can later be spatially aligned with MRI-derived measures of white-matter damage. In that setting, the model needs to support both counting discrete injured axonal profiles per unit area and estimating the total tissue area occupied by APP-positive signal. Detection rate is closely tied to the first task because it measures whether individual APP-positive components are detected as separate pathological structures rather than being missed or merged with neighboring signals.

Dice and IoU are more closely tied to the second task because they depend on how well the predicted mask reproduces the spatial extent and boundaries of the annotated signal.

This distinction helps explain why detection rate was consistently higher than Dice and IoU across training configurations and test sections. A model can correctly localize an APP-positive axonal profile even when it does not exactly match the expert-drawn boundary. This is especially relevant for small, fragmented, or low-contrast profiles, where the boundary itself may be difficult to define consistently. For this reason, detection rate provides the main readout of component-level recovery, while Dice and IoU provide complementary information about mask quality and area agreement.

The comparison across training configurations suggests that, under limited-label conditions, annotation composition may matter as much as, or more than, annotation volume. The D1+D4 configuration, which includes sections from two different animals, achieved the highest mean detection rate despite using fewer training tiles than the full D1+D2+D3+D4 configuration. In contrast, adding D2 and D3, both from P2, did not improve detection rate over the D1-only baseline. The pattern suggests that additional annotations are not equally informative. Under limited supervision, sections that broaden the training distribution may provide more useful signals than additional sections that are highly similar to data already represented in the training set. This interpretation is consistent with the broader supervised-learning principle that redundant examples can show diminishing returns and, in some cases, may contribute to overfitting [28].

One possible explanation is that adding D4 from P3 introduced a more distinct tissue appearance, injury morphology, or staining context, thereby improving the model’s exposure to inter-animal variability. However, we cannot exclude an alternative explanation: D2 and D3 may have introduced variation that was less representative of the held-out test data and therefore slightly reduced generalization. These explanations point to different but related lessons. The first supports cross-animal sampling to increase useful diversity, whereas the second cautions that not all additional annotations improve robustness if they shift the training distribution in an unhelpful direction. With the present cohort, we cannot fully distinguish between these possibilities. Because our cohort includes only three animals and four sections, we treat this finding as hypothesis-generating rather than definitive.

The paired augmentation analysis provides a related lesson about robustness. The no-spatial baseline already included histogram matching, so the comparison asked whether spatial and morphology-aware augmentation added value after inter-section stain variation had been partly controlled. Full augmentation improved performance on D2, D3, and D4 across all three metrics, while D1 showed marginal decreases. Even so, the directionally consistent gains across three of four sections suggest that spatial augmentation may provide a modest robustness benefit beyond intensity normalization, a possibility that larger annotated datasets would be better positioned to confirm.

The comparison with UniVerSeg and MicroSAM further shows why APP segmentation requires a task-specific model. MicroSAM, used without APP-specific fine-tuning, showed limited sensitivity to APP-positive axonal profiles across the evaluated tiles and produced near-zero Dice in all cases. UniVerSeg was evaluated with the largest support sets available in our dataset, but it still generated dense false-positive predictions that visually corresponded to cell nuclei and other DAB-positive background structures. These structures can share staining characteristics with injured axonal profiles, but they do not represent APP-positive axonal pathology. Separating them from true APP-positive profiles requires tissue-level contextual judgment that support-based generalization alone did not provide in this setting. PIGMENT, in contrast, was trained specifically to recognize APP-positive axonal profiles and used target-aware augmentation to model variation in morphology, staining intensity, fragmentation, and tissue context. Its advantage therefore arose not simply from selecting a different segmentation backbone, but from designing the annotation, augmentation, evaluation, and spatial reconstruction workflow around the biological properties of the target pathology. The performance gap illustrates the importance of task-specific modeling when pathological structures are sparse and visually confounded by normal tissue and staining artifacts.

The independent controlled cortical impact sample provides a qualitative test outside the RAI development cohort and illustrates a substantially different distribution of injury burden. RAI typically produces diffuse and relatively low-grade axonal pathology, whereas the CCI sample showed a larger and more spatially concentrated APP burden. PIGMENT was able to process this higher-burden section and generate a section-level spatial summary of predicted APP pathology, demonstrating the practical scalability of the workflow across qualitatively different patterns of injury. However, because exhaustive expert annotation was unavailable and some predictions occurred in regions where pathology was not expected, this experiment should not be interpreted as formal evidence of external generalization. Fully annotated external CCI and RAI sections will be needed for rigorous quantitative validation. PIGMENT may support future micros–macro association studies by making APP-positive axonal injury measurable across larger tissue regions while preserving its spatial organization. Within such studies, section-level APP predictions could be registered to ex vivo or in vivo MRI space, summarized within anatomically defined white-matter regions, and compared with local imaging abnormalities. The same maps could also be analyzed according to structural connectivity, distance from the primary injury site, or involvement of hypothesized vulnerable pathways. These analyses could test whether MRI-derived measures reflect the density, spatial extent, or network distribution of microscopic axonal injury. The present study does not validate a specific MRI biomarker; rather, it establishes the histology-side segmentation and mapping framework required before direct microscopic-to-macroscopic correspondence can be evaluated.

This study has several limitations that should guide future work. The cohort is small, with three pigs, four sections, and 525 annotated tiles, so the apparent benefit of annotation composition should be tested in larger cohorts that include more animals, staining batches, tissue regions, and injury models. The most difficult cases remain faint, fragmented, or low-contrast APP-positive profiles, where even expert labels may be locally uncertain. Future work should therefore include independently annotated external test sections or representative test regions, rather than requiring exhaustive manual annotation of entire whole-slide images. Larger external test sets would allow more rigorous evaluation of generalization, while active learning could help prioritize uncertain regions for expert review. Additional studies should evaluate whether the broader PIGMENT workflow, including limited-label training, APP-specific augmentation, component-level evaluation, and spatial reconstruction—can be adapted to other stains, tissues, and sparse histopathological targets. Such transfer should be demonstrated experimentally rather than assumed from the present APP-focused results.

In conclusion, PIGMENT provides an annotation-efficient framework for segmenting sparse and morphologically heterogeneous pathology in high-resolution histology. Although developed and evaluated for APP-positive axonal injury in porcine white matter, its core design principles, including task-specific augmentation, stain normalization, tile-based learning, spatial reconstruction, and instance-level evaluation, are applicable to a broader range of histopathology problems characterized by limited expert annotations, variable staining, fragmented structures, and highly imbalanced foreground signals. By supporting both component-level detection and pixel-level burden mapping, PIGMENT provides a scalable approach for converting local microscopic findings into spatially organized tissue-level representations. Larger and more diverse cohorts will be needed to establish external validity and to test transfer across stains, tissues, and pathological targets. Nevertheless, these results provide a practical foundation for broader computational histology applications and for future studies linking microscopic pathology with MRI-derived measures, regional tissue damage, and network-level abnormalities.

## Acknowledgments

This work was supported by the National Institutes of Health and the U.S. Department of Defense (grant numbers to be confirmed by the authors). All animal procedures were approved by the Institutional Animal Care and Use Committee (IACUC) at the University of Pennsylvania.

## Author Contributions

P.A. led the development of the PIGMENT framework, including model implementation, experiments, quantitative analyses, and manuscript preparation. J.D. contributed to the augmentation strategy, experimental design, analysis, and manuscript writing and revision. D.P. performed histology preprocessing and contributed to ground-truth preparation and validation. S.K.C.A. implemented the baseline methods, conducted qualitative comparisons, and built the containerized software environment. D.H.S. provided the porcine RAI cohort and contributed to the biological and injury-model context. H.S. provided APP annotation data and contributed to the RAI dataset and histopathological interpretation. J.A.W. contributed to study design, interpretation of the injury model, and annotation support. V.E.J. contributed to data acquisition, preprocessing, and histological interpretation. R.D.-A. contributed to interpretation of the findings in the context of traumatic brain injury. R.V. supervised the project, guided study conception and methodological development, and critically revised the manuscript. All authors reviewed and approved the final manuscript.

## Ethical Statement

All animal procedures were performed in accordance with institutional guidelines and were approved by the Institutional Animal Care and Use Committee (IACUC) at the University of Pennsylvania. All experiments complied with relevant ethical regulations for animal research.

## Informed Consent / Patient Consent

Not applicable. This study involved animal subjects only and did not include human participants.

## Trial Registration

Not applicable. This study did not involve a clinical trial.

## Grant Number

Grant numbers: R01NS123034, R01CA278819, W81XWH20-1-0838, W81XWH-20-1-0901, HT9425-23-1-1039, RD-000547.

## Supplementary Information

**Figure S1.**
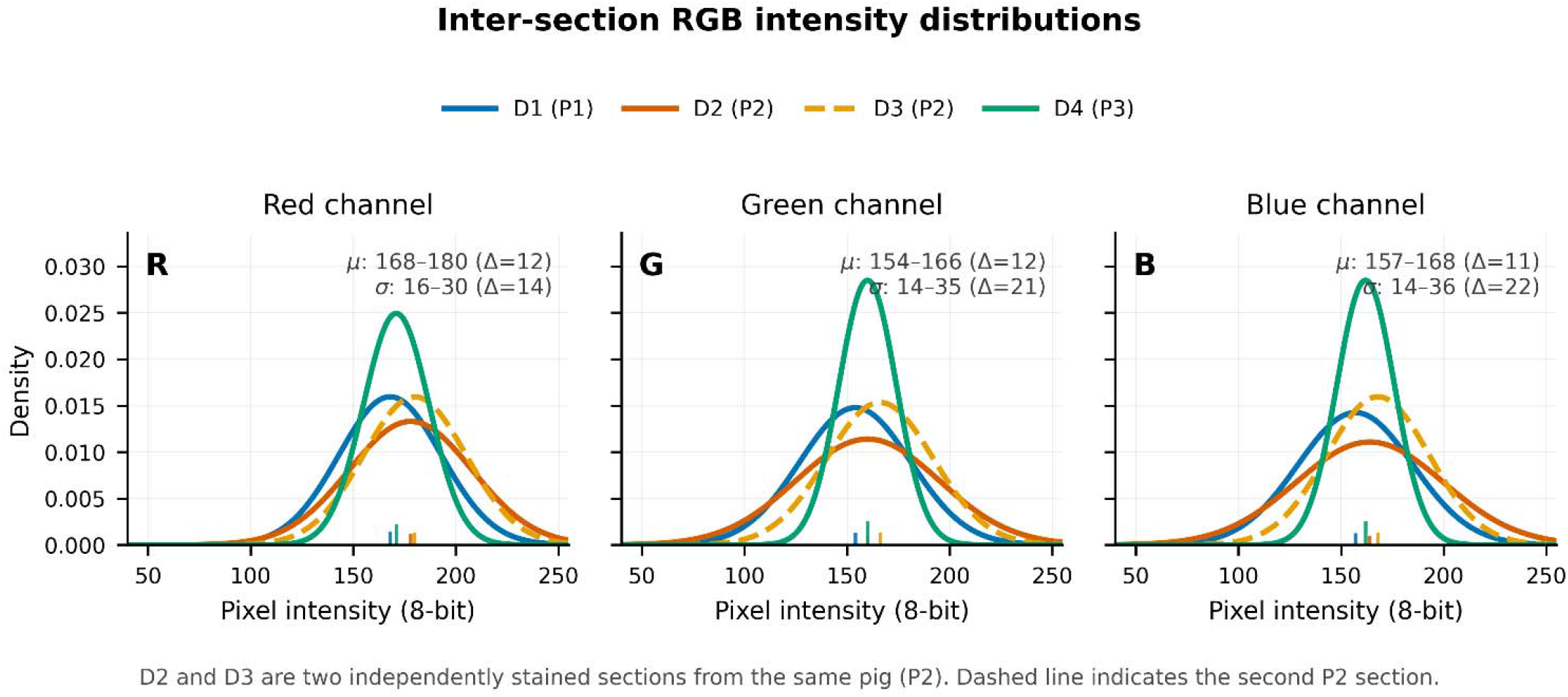
Inter-section RGB intensity distributions across APP-stained porcine white-matter sections. Gaussian approximations of 8-bit red, green, and blue channel intensity distributions are shown for the four sections used in the PIGMENT dataset. Each panel overlays the same four sections for one color channel, allowing direct comparison of staining and acquisition variability across sections. D2 and D3 are independently stained sections from the same pig, with D3 shown as a dashed line. Short vertical ticks indicate channel-specific mean intensities. Differences in curve position reflect inter-section shifts in color intensity, whereas differences in curve width reflect variation in within-section intensity dispersion. These batch-level differences motivate the photometric augmentation strategy used during PIGMENT training.

**Figure S2.**
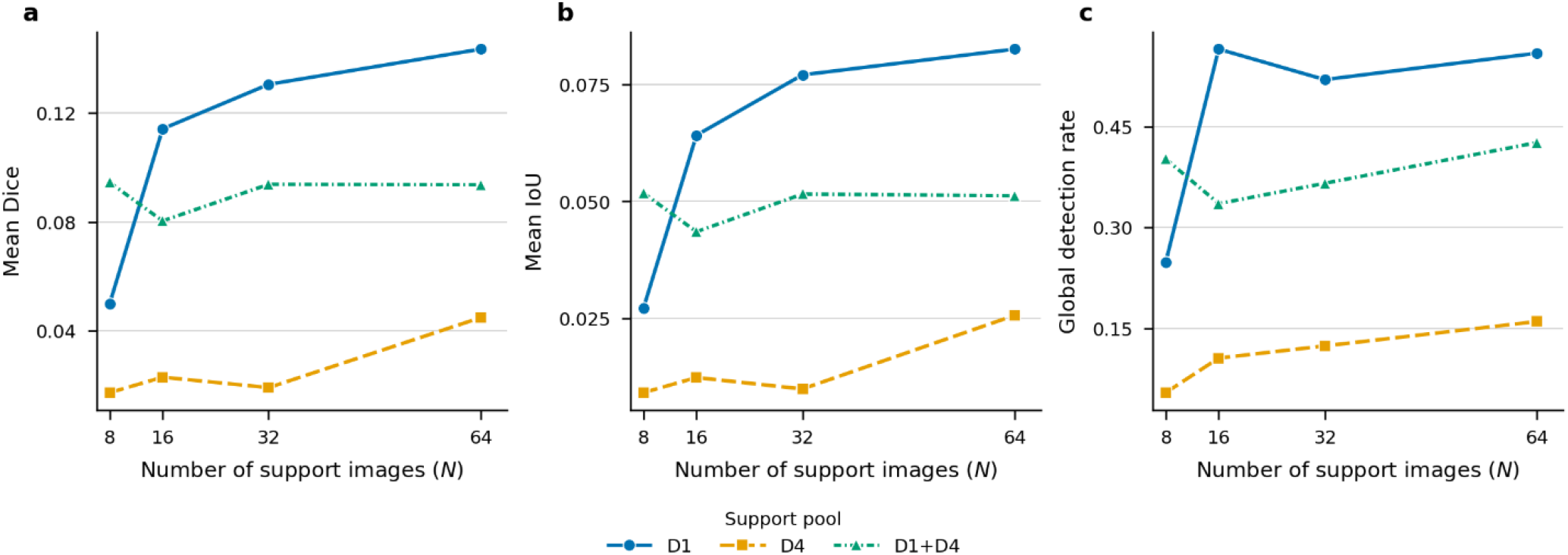
Dice, IoU, and DR for the PIGMENT model with geometric augmentations across the four training configurations. The pattern matches Fig. 4 (full pipeline): DR is consistently higher than pixel-overlap metrics, and the D1+D4 configuration is strongest overall.

**Figure S3.**
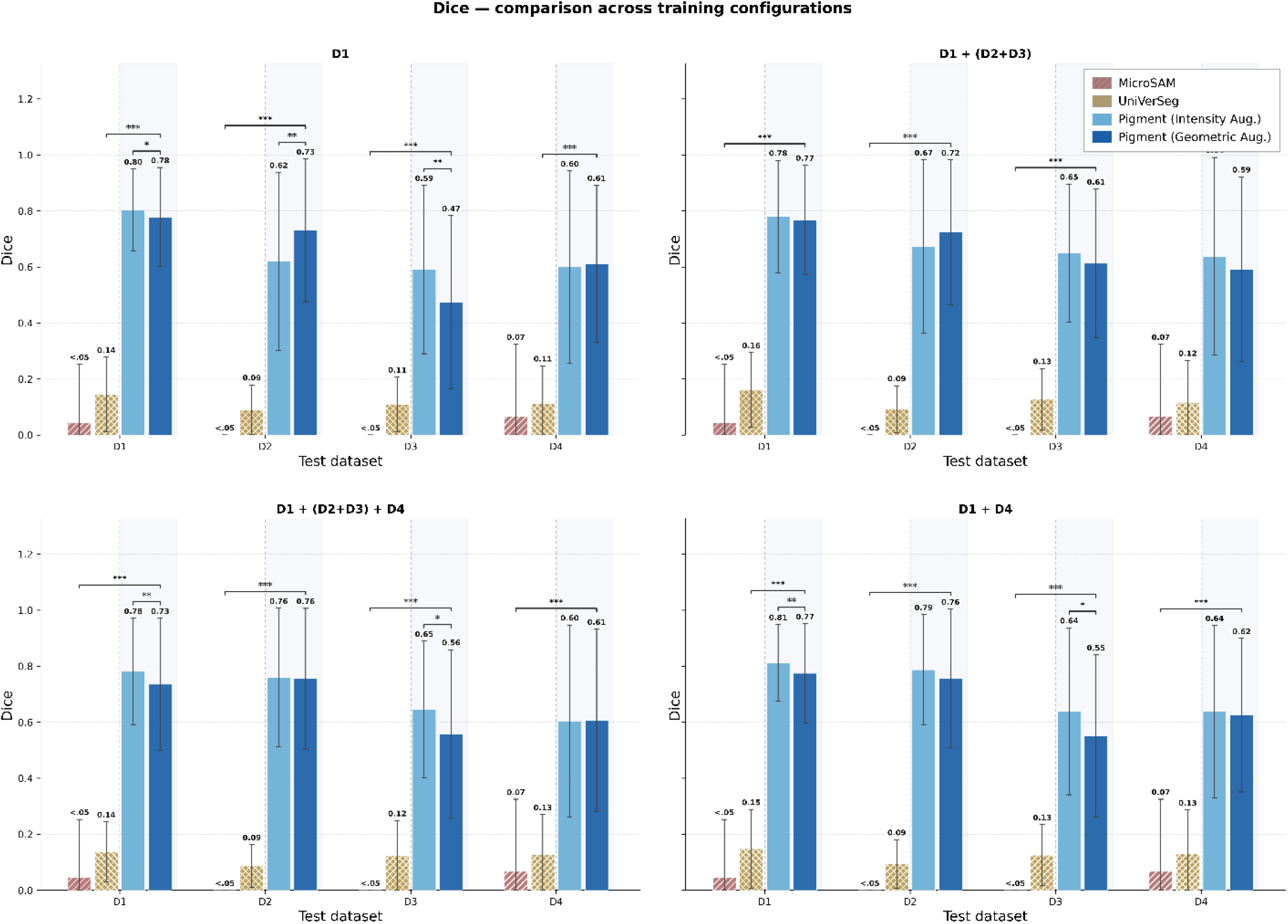
Dice across test sections and training configurations for MicroSAM, UniverSeg, and PIGMENT under intensity-only and full geometric augmentation. PIGMENT achieves the highest Dice in most settings.

**Figure S4.**
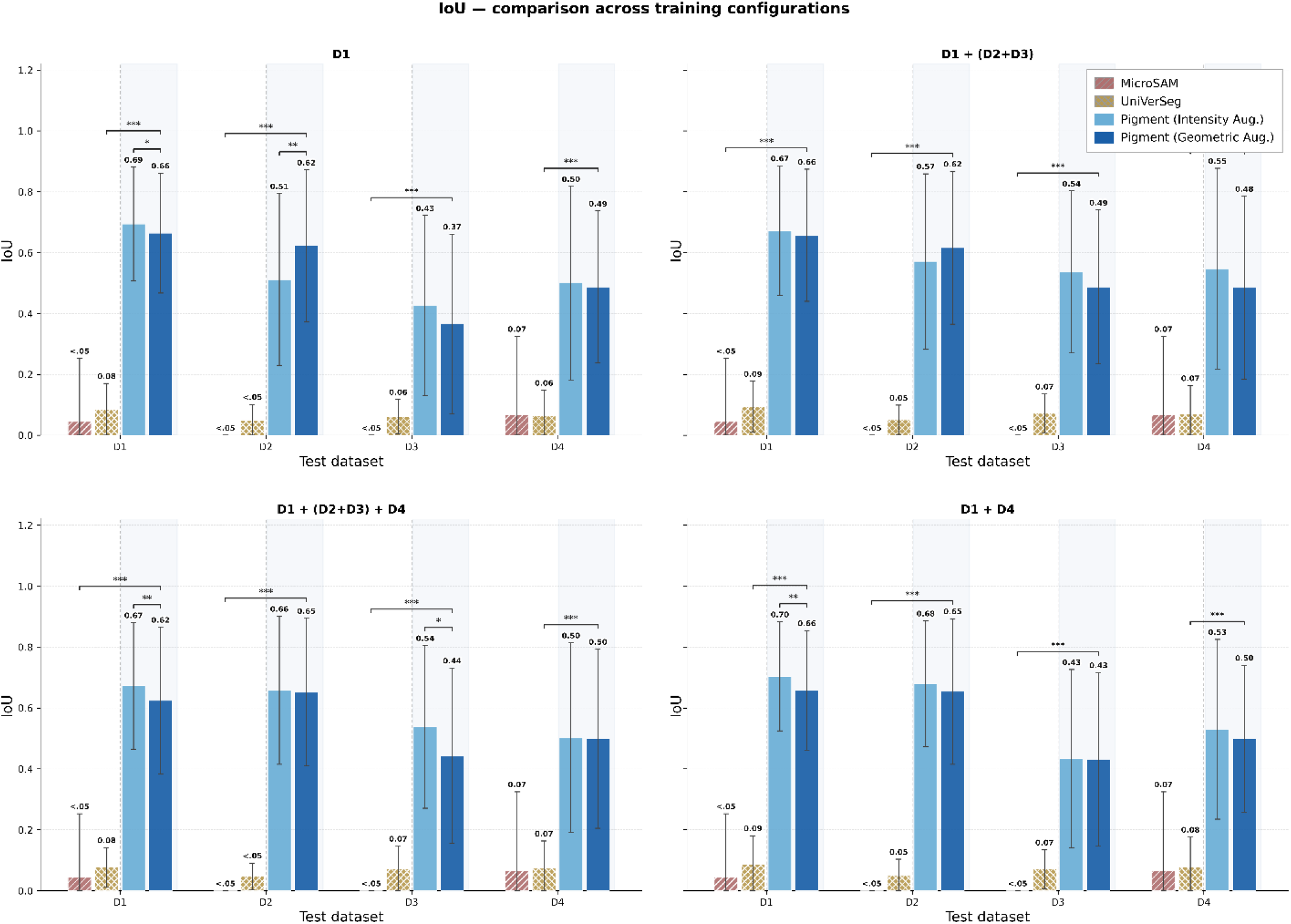
Intersection-over-union (IoU) across test sections and training configurations for MicroSAM, UniverSeg, and PIGMENT. PIGMENT achieves the highest IoU in most settings, consistent with the Dice and DR comparisons.

**Figure S5.**
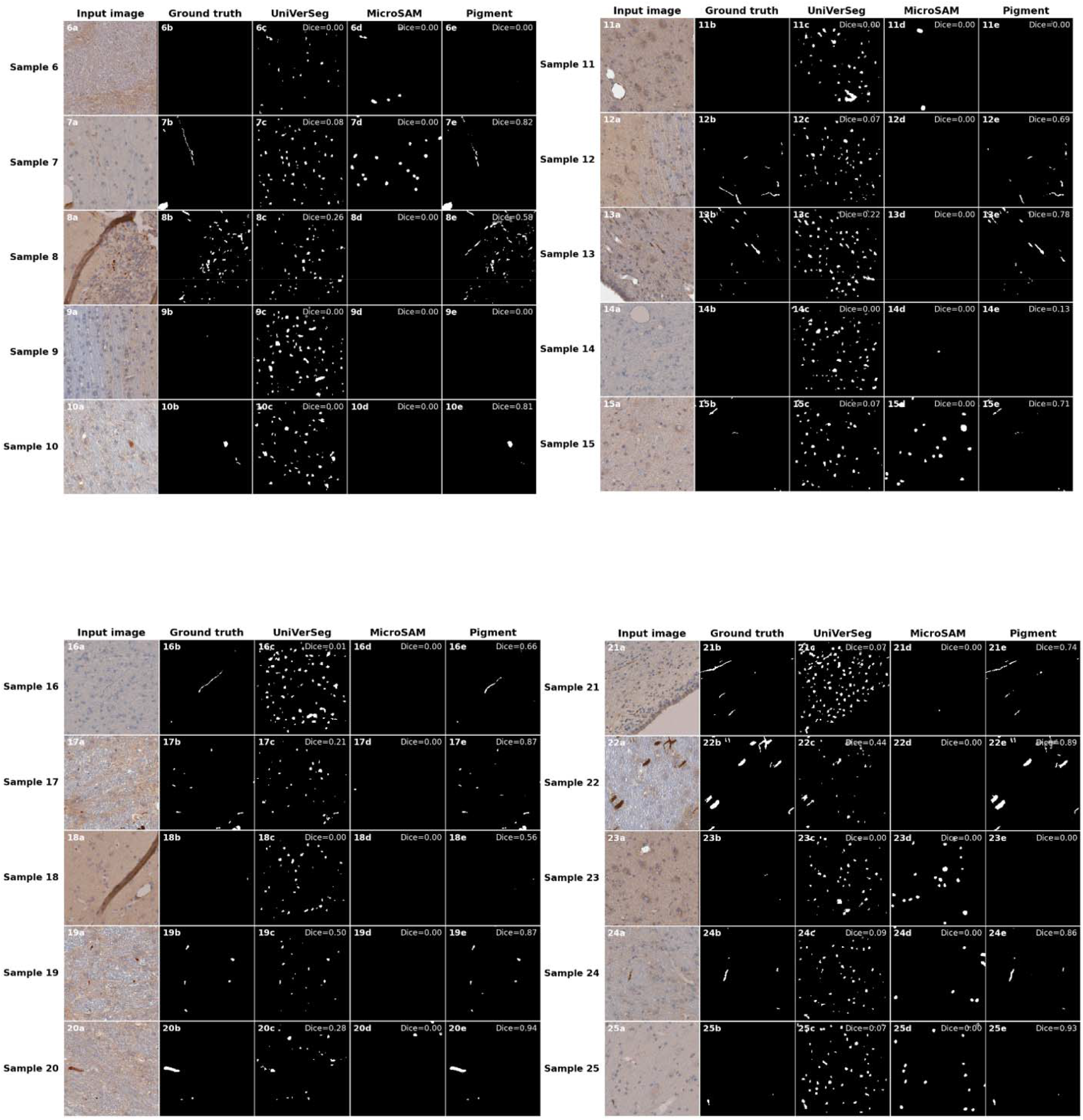
Extended qualitative comparison across diverse APP morphologies, staining conditions, and annotation regimes (Samples 6-15). Representative held-out histology tiles complementing Fig. 5. Each row shows input tile, expert ground-truth annotation, UniverSeg prediction, MicroSAM prediction, and PIGMENT prediction. These supplementary samples span APP-negative windows, sparse isolated deposits, elongated axonal fibers, fragmented annotations, dense heterogeneous APP regions, and low-contrast cases. UniverSeg over-segments background tissue with fragmented false positives, and MicroSAM under-segments or fails entirely in sparse and elongated morphologies.

### Mathematical formulations

This section provides the complete mathematical specification of the components summarized in the main text Materials and Methods: the SegFormer-B0 encoder–decoder, the training objective and EMA teacher regularization, the target-aware augmentation transforms, and the post-processing applied before evaluation. Equations are numbered S1, S2, … for ease of reference.

### Architecture details

The segmentation model is a function fθ: ℝ □×W×^3^ → [0, 1] □ ×W that maps an RGB tile of size H × W to a pixel-wise probability of APP positivity. We instantiate fθ with SegFormer-B0, a hierarchical vision transformer for semantic segmentation. The encoder (MiT-B0) is composed of four stages; at each stage, features F□ are produced by an encoder block E□ applied to the previous feature map, with spatial resolution progressively halved and channel dimensionality increased (C1 = 32, C2 = 64, C3 = 160, C4 = 256):

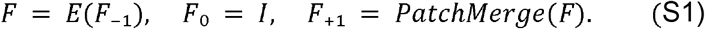

Query, key, and value matrices are obtained via linear projections of the input feature matrix X with learnable projection matrices W_Q, W_K, W_V:

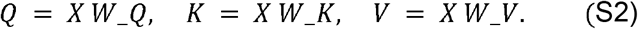

Self-attention at each stage uses Efficient Self-Attention with a sequence-reduction ratio R□, reducing the effective key–value length from N = H□W□ to N / R□ and lowering attention complexity from O(N^2^) to O(N^2^ / R□):

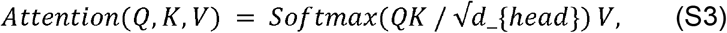

where d_{head} is the head dimension. Positional information is injected via Mix-FFN blocks that combine a depthwise 3 × 3 convolution with two linear projections and a GELU non-linearity:

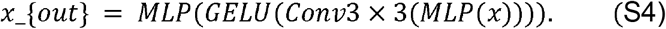

The All-MLP decoder upsamples each encoder feature map F□ to a common spatial resolution, channel-concatenates the four upsampled maps, projects them with a linear layer, and bilinearly upsamples the logits to the input resolution:

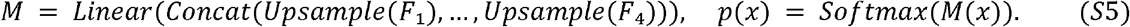

The model contains approximately 3.7 million parameters. The multi-section training strategy is architecture-agnostic and can be combined with any encoder–decoder segmentation network.

### Training objective and EMA teacher

Let V = {x : L(x) ≠ 255} denote the set of valid pixels, that is, pixels not marked as ignore. The training objective is pixel-wise cross-entropy computed only over V:

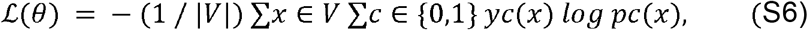

where yc(x) ∈ {0, 1} is the one-hot ground-truth label for class c and pc(x) is the predicted probability from a channel-wise Softmax on the upsampled logits. No explicit class weighting is applied; class imbalance is addressed through the augmentation pipeline.

A teacher network with parameters θ□⍰ is maintained as an exponential moving average of the student parameters □⍰ with decay d = 0.999. The teacher receives no gradient updates and is refreshed at every step according to

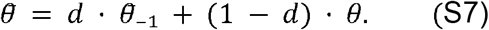

The teacher flags pixel x as ambiguous (and excludes it from the gradient for that step) when its prediction strongly disagrees with the manual label in a high-confidence region:

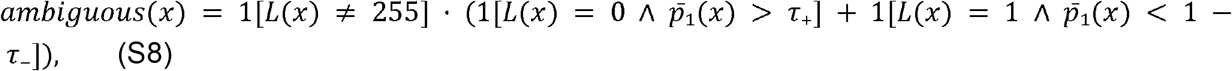

with τ_+_ = τ_−_ = 0.97 and p□_1_(x) the teacher’s class-1 (APP-positive) probability at pixel x. Pixels flagged ambiguous in a given step are temporarily assigned the ignore value 255 and excluded from the cross-entropy loss for that step. This is a Mean-Teacher-style regularization that reduces the influence of likely mis-annotated pixels (faint deposits missed by the annotator, boundary pixels of small fragmented components, or staining artifacts inadvertently included in foreground).

Models are optimized with AdamW (learning rate η = 1 × 10^−^□, β_1_ = 0.9, β_2_ = 0.999, weight decay λ = 0.01), batch size 8, with dropout applied to encoder hidden states (p = 0.1), attention weights (p = 0.1), and the classifier head (p = 0.3).

### Augmentation transforms (formal definitions)

Data augmentation is formulated as a stochastic transformation T sampled from a distribution P (τ) that acts jointly on the image– mask pair (I, L):

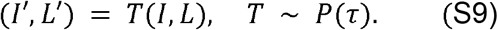

The model is trained to minimize the expected cross-entropy loss over augmented samples, where D is the dataset distribution:

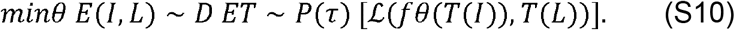

For copy-paste operations, APP-positive components are sampled only from the ground-truth mask of the same training image, so that no labels are transferred across images. Spatial transformations act jointly on image and mask to preserve pixel-wise correspondence; photometric transformations act on the image only. Pixels assigned the ignore label 255 are preserved through all transforms and excluded from loss computation. Below, x denotes a 2D spatial coordinate inside the tile and D the tile width.

#### Affine transform

Each spatial location x is mapped according to

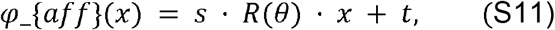

where R(θ) is a 2×2 rotation matrix with θ ~ U (−18°, 18°), scale s ~ U(0.96, 1.06), and translation t_{x}, t_{y} ~ U (−0.06 D, 0.06 D). Image pixels are resampled with bilinear interpolation; masks with nearest-neighbour interpolation.

#### Global RGB shift

Each colour channel c ∈ {R, G, B} is shifted by an additive offset

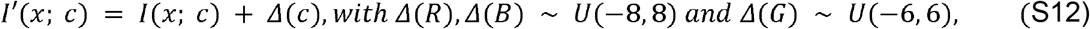

in the [0, 255] pixel range. The narrower green-channel range reflects its greater sensitivity to APP immunoreactivity in brightfield histology.

#### Multi-instance copy-paste

Let C ⍰ denote the k-th sampled connected component from L. Each instance is rigidly transformed,

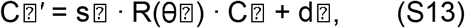

with n ~ U({5, …, 20}) instances per image, scale s⍰ ~ U(0.90, 1.25), in-plane rotation θ⍰ ~ U(−8°, 8°), and placement offset d⍰ sampled uniformly over the tile. An optional elastic deformation with amplitude α ~ U(0, 180) and smoothness σ ~ U(0, 5) is applied to introduce sub-pixel shape variation, and a 2-pixel feather blend is applied at the component boundary to suppress visible seams.

#### GMM-guided label promotion

A Gaussian Mixture Model with K = 4 components is fitted to the RGB distribution of manually annotated APP-positive pixels. For a pixel x with colour vector I(x), the likelihood under the mixture is

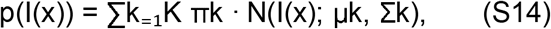

and the posterior probability that pixel x belongs to mixture component k is

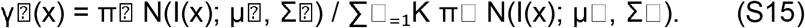

Pixels whose likelihood exceeds a data-driven threshold and that lie within a bounded ROI (at most 224× 224 pixels, at most 3 candidate regions, with no more than 4500 promoted pixels per image) are promoted to foreground. The strict spatial and count constraints prevent promotion of false positives in regions of clearly negative tissue.

#### Ground-truth neighborhood diffusion

The binary mask L is convolved with a Gaussian kernel of radius r ~ U(4, 10) pixels to produce a soft proximity map L. A blended image Ĩ is formed as

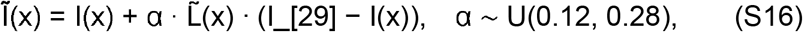

where I_[29] is the mean intensity of APP-positive pixels. The blend is gated by the condition L(x) ≥ T, with T ~ U(0.75, 0.90), restricting the modification to regions immediately adjacent to annotated foreground and preventing label spread into clearly negative tissue. A complete listing of transform categories and hyperparameters is given in Table S2.

### Post-processing and evaluation metrics (formal definitions)

#### Morphological post-processing of predictions

For each connected component C⍰ in the predicted binary mask, the minimum bounding rectangle yields principal-axis dimensions (w⍰, h⍰). The elongation ratio is

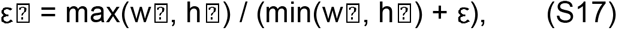

with ε = 10^−^□ for numerical stability. Components satisfying both area(C⍰) ≥ T_{area} and εIZ ≥ τ_{ε} are suppressed (set to background); we use τ_{area} = 2300 pixels and τ_
{ε} = 3.0. Thresholds were chosen so that no annotated APP-positive component in the training set is excluded, and structures of this scale and shape correspond to staining artifacts at fold edges or tissue tears rather than APP-positive profiles.

#### Boundary-aware ignore masking

For each connected component C⍰ ^{GT} in the binary ground-truth mask, the component is excluded from evaluation if it both (i) intersects the patch border, i.e. there exists x ∈ C⍰ ^{GT} with x ∈ ∂Ω, and (ii) has area below a threshold, area(C⍰ ^{GT}) < τ_{edge} with τ_{edge} = 50 pixels. The resulting ignore mask M_[30] is applied to both ground truth and prediction before metric computation:

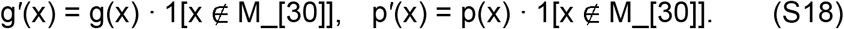

#### Detection rate construction

Small gaps within the GT foreground are first closed using morphological closing with a disk-shaped structuring element B□ of radius r = 2:

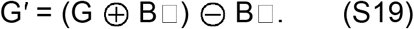

The predicted mask is dilated by B□ ′ with r′ = 1 to introduce a small spatial tolerance, P′ = P ⊕ B□ ′, and predicted components with area below τ⍰ □ = 25 pixels are discarded. A ground-truth component C⍰ ^{GT} is considered valid if area (C⍰ ^{GT}) ≥ τ□□ = 25 pixels and it has not been excluded by the boundary-aware ignore mask. A valid component is detected if

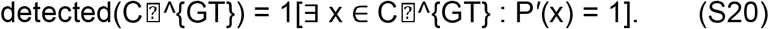

The global detection rate (Eq. 2 in the main text) is computed by summing detections over all test tiles and dividing by the total number of valid components; here we have reproduced the construction in full to make the relationship between the morphological tolerances (r, r′) and the metric explicit.

### Ground-truth binarization

The annotator’s overlay was rendered as a distinct green channel; after luminance conversion to grayscale, the annotation pixels were separated from background tissue by an intensity threshold (pixels with grayscale intensity at least 228 were assigned APP-positive, pixels at most 227 were assigned background). The threshold was determined empirically on representative training tiles and verified to reproduce the annotator’s intent. PIGMENT consumes the resulting binary masks directly.

